# Evolution and expansion of multidrug resistant malaria in Southeast Asia: a genomic epidemiology study

**DOI:** 10.1101/621763

**Authors:** William L Hamilton, Roberto Amato, Rob W van der Pluijm, Christopher G Jacob, Huynh Hong Quang, Nguyen Thanh Thuy-Nhien, Tran Tinh Hien, Bouasy Hongvanthong, Keobouphaphone Chindavongsa, Mayfong Mayxay, Huy Rekol, Rithea Leang, Cheah Huch, Lek Dysoley, Chanaki Amaratunga, Seila Suon, Rick M Fairhurst, Rupam Tripura, Thomas J Peto, Yok Sovann, Podjanee Jittamala, Borimas Hanboonkunupakarn, Sasithon Pukrittayakamee, Nguyen Hoang Chau, Mallika Imwong, Mehul Dhorda, Ranitha Vongpromek, Xin Hui S Chan, Richard J Maude, Richard D Pearson, T Nguyen, Kirk Rockett, Eleanor Drury, Sonia Gonçalves, Nicholas J White, Nicholas P Day, Dominic P Kwiatkowski, Arjen M Dondorp, Olivo Miotto

## Abstract

**Background:** A multidrug resistant co-lineage of *Plasmodium falciparum* malaria, named KEL1/PLA1, spread across Cambodia c.2008-2013, causing high treatment failure rates to the frontline combination therapy dihydroartemisinin-piperaquine. Here, we report on the evolution and spread of KEL1/PLA1 in subsequent years.

**Methods:** We analysed whole genome sequencing data from 1,673 *P. falciparum* clinical samples collected in 2008-2018 from northeast Thailand, Laos, Cambodia and Vietnam. By investigating genome-wide relatedness between parasites, we inferred patterns of shared ancestry in the KEL1/PLA1 population.

**Findings:** KEL1/PLA1 spread rapidly from 2015 into all of the surveyed countries and now exceeds 80% of the *P. falciparum* population in several regions. These parasites maintained a high level of genetic relatedness reflecting their common origin. However, several genetic subgroups have recently emerged within this co-lineage with diverse geographical distributions. Some of these emerging KEL1/PLA1 subgroups carry recent mutations in the *chloroquine resistance transporter* (*crt*) gene, which arise on a specific genetic background comprising multiple genomic regions.

**Interpretation:** After emerging and circulating for several years within Cambodia, the *P. falciparum* KEL1/PLA1 co-lineage diversified into multiple subgroups and acquired new genetic features including novel crt mutations. These subgroups have rapidly spread into neighbouring countries, suggesting enhanced fitness. These findings highlight the urgent need for elimination of this increasingly drug-resistant parasite co-lineage, and the importance of genetic surveillance in accelerating elimination efforts.

**Funding:** Wellcome Trust, Bill & Melinda Gates Foundation, UK Medical Research Council, UK Department for International Development.

**Research in context:** ##### Evidence before this study

This study updates our previous work describing the emergence and spread of a multidrug resistant *P. falciparum* co-lineage (KEL1/PLA1) within Cambodia up to 2013. Since then, a regional genetic surveillance project, GenRe-Mekong, has reported that markers of dihydroartemisinin-piperaquine (DHA-PPQ) resistance have increased in frequency in neighbouring countries. A PubMed search (terms: “artemisinin”, “piperaquine”, “resistance”, “southeast asia”) for articles listed since our previous study (from 30/10/2017 to 05/01/2019) yielded 28 results, including reports of a recent sharp decline in DHA-PPQ clinical efficacy in Vietnam; the spread of genetic markers of DHA-PPQ resistance into neighbouring countries by Imwong and colleagues; and multiple reports associating mutations in the *crt* gene with piperaquine resistance, including newly emerging *crt* variants in Southeast Asia.

##### Added value of this study

We analysed *P. falciparum* whole genomes collected up to early 2018 from Eastern Southeast Asia (Cambodia and surrounding regions), describing the fine-scale epidemiology of multiple KEL1/PLA1 genetic subgroups that have spread out from Cambodia since 2015 and taken over indigenous parasite populations in northeastern Thailand, southern and central Vietnam and parts of southern Laos. Several newly emerging *crt* mutations accompanied the spread and expansion of KEL1/PLA1 subgroups, suggesting an active proliferation of biologically fit, multidrug resistant parasites.

##### Implications of all the available evidence

The problem of *P. falciparum* multidrug resistance has dramatically worsened in Eastern Southeast Asia since previous reports. KEL1/PLA1 has diversified and spread widely across Eastern Southeast Asia since 2015, becoming the predominant parasite group in several regions. This may have been fuelled by continued parasite exposure to DHA-PPQ, resulting in sustained selection after KEL1/PLA1 became established. Continued drug pressure enabled the acquisition of further mutations, resulting in higher levels of resistance. These data demonstrate the value of pathogen genetic surveillance and the urgent need to eliminate these dangerous parasites.

## Introduction

In recent years, frontline treatments for *Plasmodium falciparum* malaria have been failing in parts of Southeast Asia^1–3^, a historic epicentre for the emergence and spread of antimalarial drug resistance^4^. The current treatment for *P. falciparum* consists of a fast-acting artemisinin derivative and a longer-acting partner drug, termed artemisinin combination therapy (ACT). Dihydroartemisinin with piperaquine (DHA-PPQ) has been the ACT of choice in Cambodia, Vietnam and Thailand for lengthy periods during the last decade. Around 2008, parasites in western Cambodia began developing DHA-PPQ resistance, manifest first through delayed clearance in response to artemisinins (now widespread in Southeast Asia^5–9^), and later with the addition of resistance to the partner drug piperaquine^1,3,8^. By 2013, DHA-PPQ failed to clear *P. falciparum* infections in 46% of patients treated in western Cambodia^3^. This arose on a background of extensive pre-existing resistance to multiple antimalarials, leaving limited treatment options and threatening the future of malaria control and elimination in the region.

Large-scale genetic analyses have revealed the detailed epidemiology of drug resistance^10–15^, complementing the clinical observation of increasing rates of treatment failure. Non-synonymous mutations in *kelch13* – the most prevalent of which is the C580Y mutation^9,11,16^ – and amplification of *plasmepsin*-*II* and *plasmepsin*-*III*^17–20^ have proved valuable markers for tracking artemisinin and piperaquine resistance, respectively. These genetic markers increased in frequency across the eastern part of the Greater Mekong Subregion from 2008 to 2015^12–15^, corresponding with the spread of DHA-PPQ treatment failure. Whole genome sequencing has provided an even deeper insight into the movement, demographics and evolution of parasites, addressing questions about the origins and mechanisms of spread of these mutations^12,13^. In 2018, detailed analyses of a large whole genome dataset, including samples up to 2013, revealed that most parasites with the *kelch13* C580Y mutation and the *plasmepsin*-*II*/*III* amplification were derived from a single parasite co-lineage, termed KEL1/PLA1, that arose in western Cambodia^13^.

Such analyses raised several key uncertainties surrounding the future of KEL1/PLA1. Would these parasites continue their aggressive spread out from Cambodia? Would they spread clonally or heterogeneously? Could they evolve even higher levels of drug resistance or increased fitness? Recently, it was reported that newly emerging mutations in the *chloroquine resistance transporter* (*crt*) gene caused piperaquine resistance *in vitro*^21–23^. These *crt* substitutions occurred on a *plasmepsin*-*II*/*III* amplified genetic background, raising the question of how mutations at multiple loci interact to yield DHA-PPQ resistant phenotypes; where and by what process the new *crt* mutations are spreading; and how these mutations relate to the evolution and expansion of the KEL1/PLA1 co-lineage. To address these questions, we investigated the genomic epidemiology of DHA-PPQ resistant parasites using the most recent *P. falciparum* genomic dataset currently available, including samples up to early 2018, collected across the region through the MalariaGEN *P. falciparum* Community Project.

## Results

### Extensive international expansion and invasion of genetically related drug resistant parasites

We analysed a dataset of 2,443 whole parasite genomes collected in the period 2007-2018 from northeast Thailand, Cambodia, Laos and Vietnam, which we refer to as Eastern Southeast Asia (ESEA), contributed to the MalariaGEN *P. falciparum* Community Project. We focused on ESEA because KEL1/PLA1 has not been found outside of this region to date, and because parasites from ESEA are genetically separated from those found further west, because of a malaria-free geographic corridor running through central Thailand^12^. After removing replicates, samples with low coverage and highly diverse infections (*F*_*WS*_ < 0.95) we analysed a dataset of 1,673 samples, of which the KEL1/PLA1 status could be reliably determined in 1,615 (Supplementary Table 1, Supplementary Figure 1). We genotyped each of these samples at 56,676 well-covered single nucleotide polymorphism (SNP) variants.

Within this sample set, we identified 996 *kelch13* mutant parasites, of which 816 (82%) were C580Y. The vast majority of these C580Y mutants (802/816, 98%) belonged to the KEL1 lineage, denoting a specific haplotype surrounding the *kelch13* locus and a single epidemiological origin in western Cambodia^13^. Of the KEL1 parasites, 551/802 (69%) carried an amplification of the *plasmepsin*-*II*/*III* genes with a shared surrounding haplotype, here named PLA1, also consistent with a single epidemiological origin at this locus (Supplementary Figure 2). The tendency for KEL1 to co-occur with PLA1 increased significantly from 2007-2011 to 2016-2018 (*r*^*2*^ 0.28 vs 0.41, respectively), and the majority of parasites sampled in later years were KEL1/PLA1 (354/695, 51%), reflecting the continued expansion of this co-lineage (Figure 1A). Before 2009, KEL1/PLA1 was only found in western Cambodia; by 2016-2018 its prevalence had risen to >50% in all regions sampled except Laos (Figure 1B). This rapid rise was particularly dramatic in northeast Thailand and Vietnam, where over 80% of recent samples were KEL1/PLA1, despite their absence from these areas just eight years earlier. This is consistent with near-wholescale replacement of indigenous parasite populations with KEL1/PLA1 in these areas.

**Figure 1.**
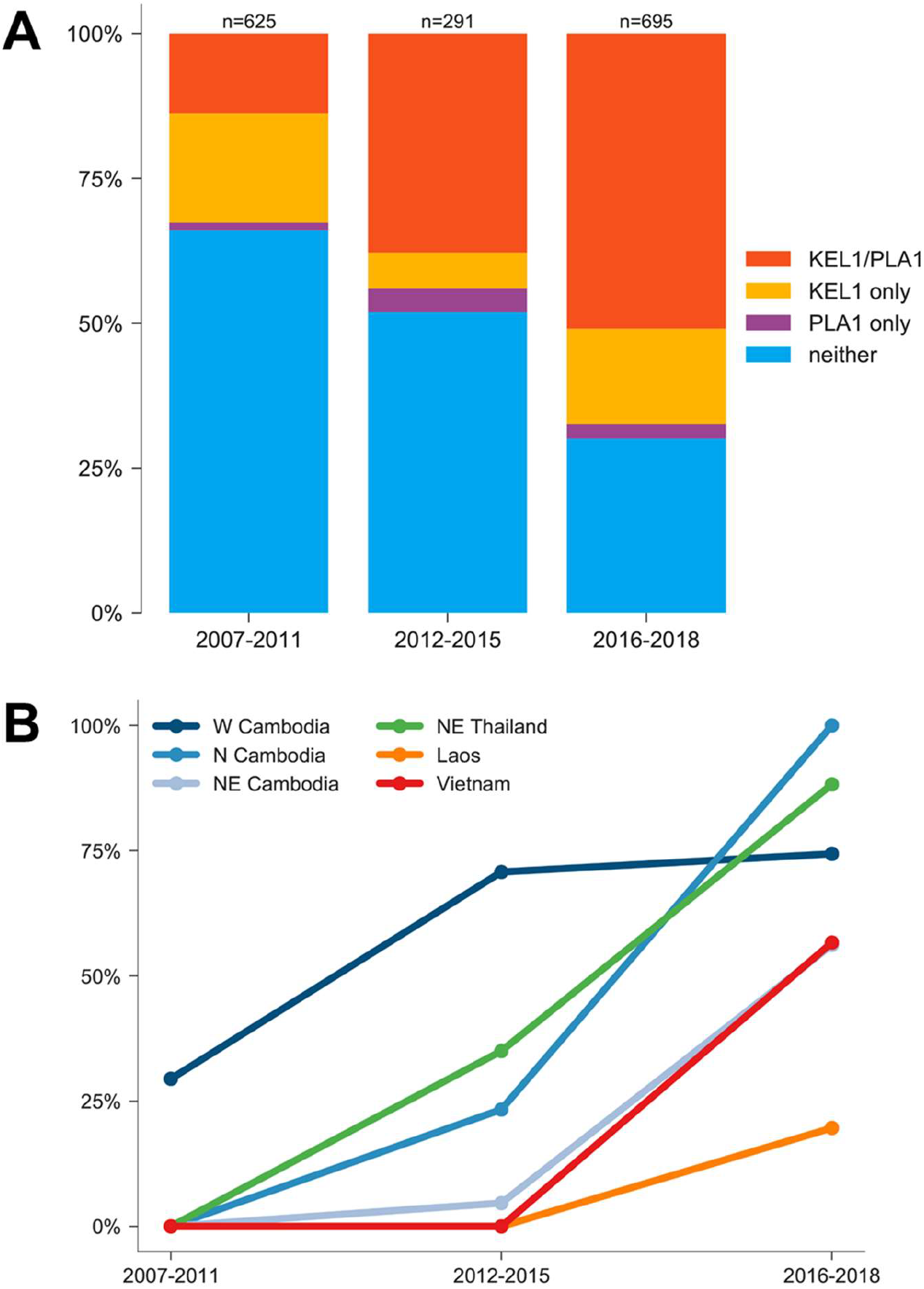
Rise in KEL1/PLA1 prevalence over tiem in ESEA countries. (A) Proportions of different combinations of *kelch13* and *plasmepsin*-*II*/*III* alleles, over three periods (2007-2011, 2012-2015 and 2016-2018) in the ESEA regions surveyed in this study. (B) Change in the frequency of KEL1/PLA1 parasites over the same periods in different ESEA geographic regions.

In previous work, we demonstrated that KEL1/PLA1 parasites from northeast Cambodia were genetically similar to those from western Cambodia, consistent with direct spread from western to northeast Cambodia^13^. We extended this analysis by examining the pattern of genetic similarity between parasites across the entire ESEA region. Overall, KEL1/PLA1 parasites had lower genetic diversity than non-KEL1/PLA1 parasites (median = 0.032 and 0.073, respectively; *P*<10^−16^, Mann-Whitney *U* test) (Figure 2A, Supplementary Figure 3). Importantly, they were genetically more similar to each other than to non-KEL1/PLA1 parasites regardless of their geographic origins; e.g. KEL1/PLA1 parasites from Vietnam were more similar to KEL1/PLA1 parasites from the rest of ESEA than to other types of Vietnamese parasites (*P*<10^−16^ for all comparisons, Mann-Whitney *U* test) (Figure 2B, Supplementary Figure 4). This is consistent with the view that KEL1/PLA1 is an invading population of western Cambodian origins spreading into surrounding countries.

**Figure 2.**
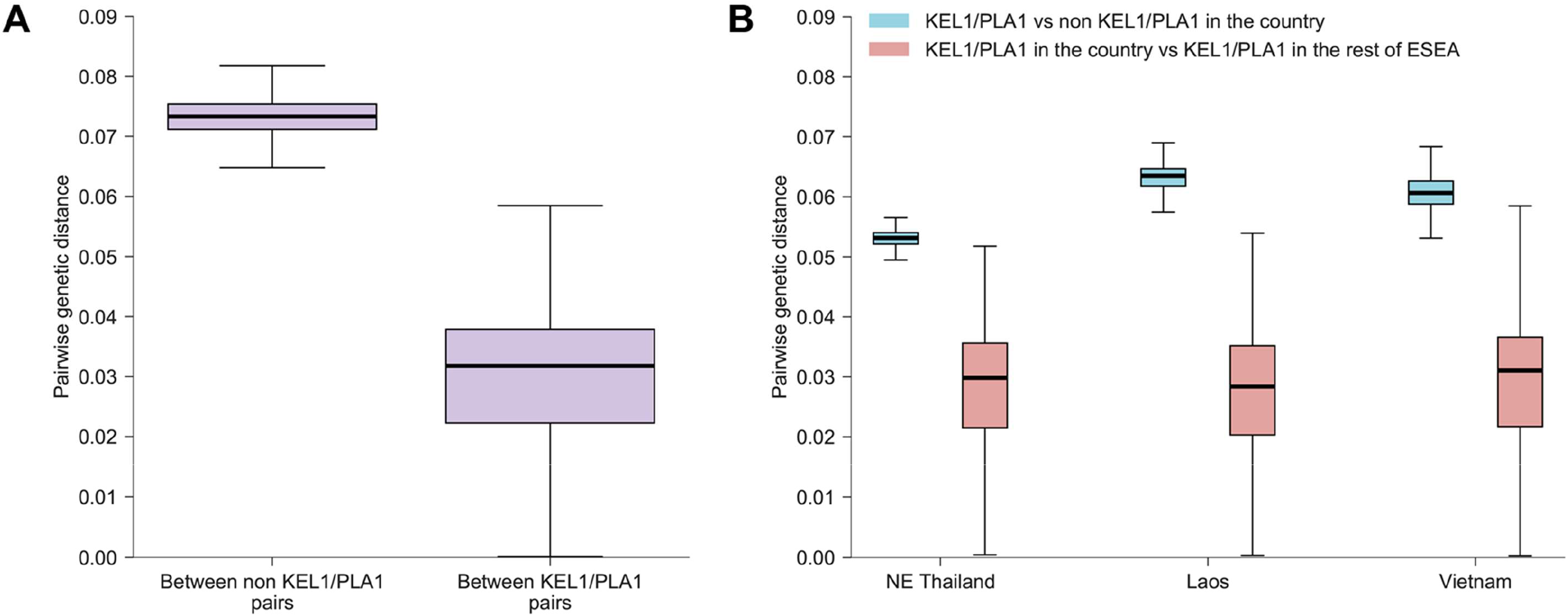
Genetic similarity aom ngst KEL1/PLA1 parasites across geographic regions. (A) Boxplot comparing the distribution of pairwise genetic distance in parasites carrying neither KEL1 nor PLA1 haplotypes, with the distribution in KEL1/PLA1 parasites. (B) Boxplot comparing pairwise distance distribution in KEL1/PLA1 parasites (red) against that in wild-type parasites (blue) from the same geographical regions.

### Diversity within the KEL1/PLA1 family

Given the high degree of genetic similarity between KEL1/PLA1 parasites, we investigated whether they have spread through a single clonal expansion or as multiple independent subgroups. Hierarchical clustering of pairwise genetic distances was used to identify groups of more closely related KEL1/PLA1 parasites (Figure 3). We defined *subgroups* of related parasites as those whose pairwise genetic distance was in the lower quartile of the KEL1/PLA1 population (Supplementary Figure 5), and numbered these *subgroups* by order of size. The six largest subgroups together comprised >50% of KEL1/PLA1 samples, and broadly captured the largest expansions of near-identical parasites, with low genetic diversity within each subgroup (Supplementary Figure 6). The subgroups had distinct geographic, temporal and genetic properties, reflecting separate epidemiological and evolutionary histories (Supplementary Figure 7, Supplementary Table 2).

**Figure 3.**
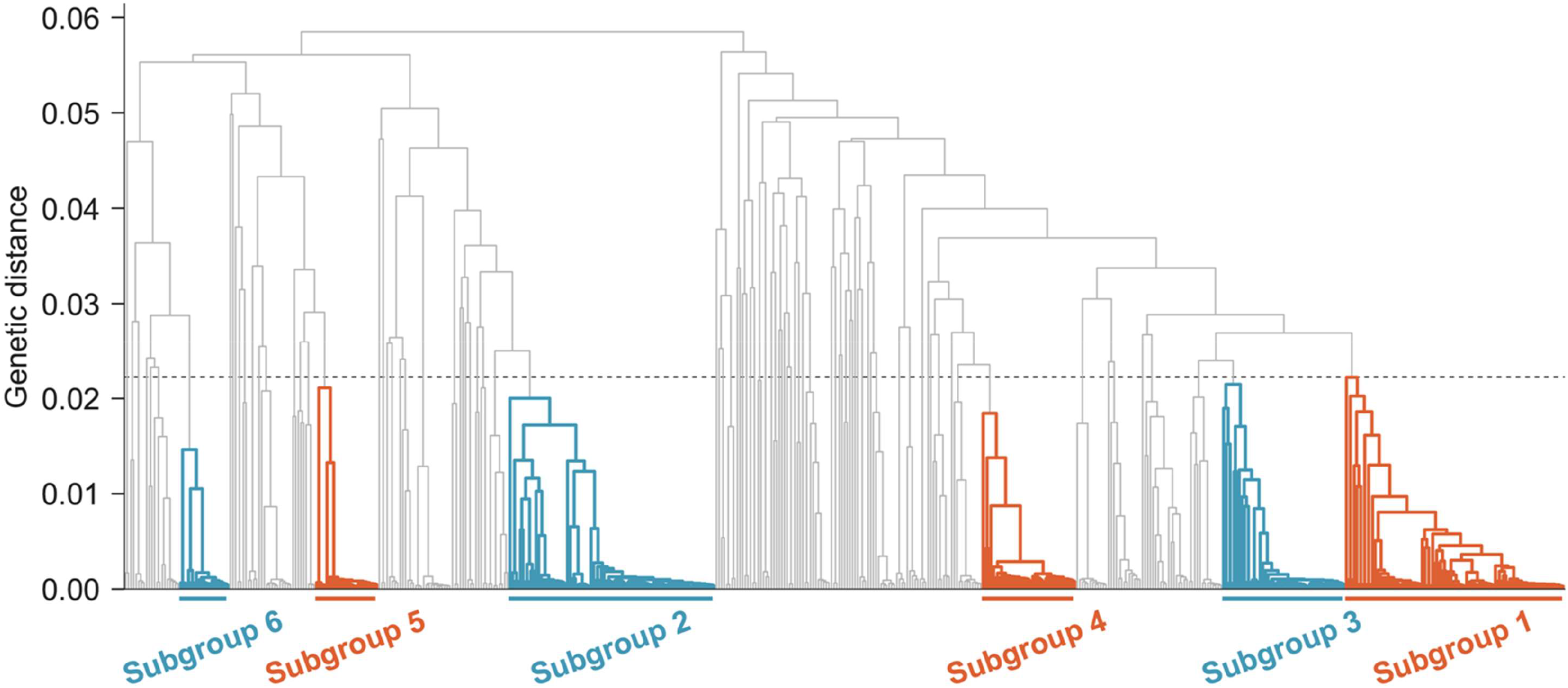
KEL1/PLA1 family tree. The dendrogram shows a hierarchical clustering tree of genetic distances within KEL1/PLA1 samples across ESEA; longer branches indicate more distant relationships. The six largest subgroups (n≥20 samples) of highly related parasites are shown in colour, and labelled below the tree. These subgroups, numbered in order of decreasing size, were identified by grouping samples with pairwise genetic distances in the lowest quartile (delimited by a dotted line).

Subgroups 1 (n= 84), 2 (n= 79) and 3 (n= 47) mostly emerged since 2016 and were all present in Cambodia, Vietnam and Laos (Figure 4A-B). Of note, these larger KEL1/PLA1 subgroups were not geographically restricted and co-existed simultaneously in the same locations across ESEA. This combination of high genetic similarity and broad geographic dispersal over just a few years implies rapid proliferation and expansion in independent overlapping transmission waves, suggesting they possess a selective advantage. In contrast, subgroup 4 (n=36) and subgroup 6 (n=19) were largely confined to Cambodia; these subgroups were responsible for the initial KEL1/PLA1 expansion in 2007-2011, but subsequently became uncommon. We also identified smaller subgroups with limited geographic and temporal distributions, such as subgroup 5 (n=24) which was almost exclusively found in northeast Cambodia in 2016-2017.

**Figure 4.**
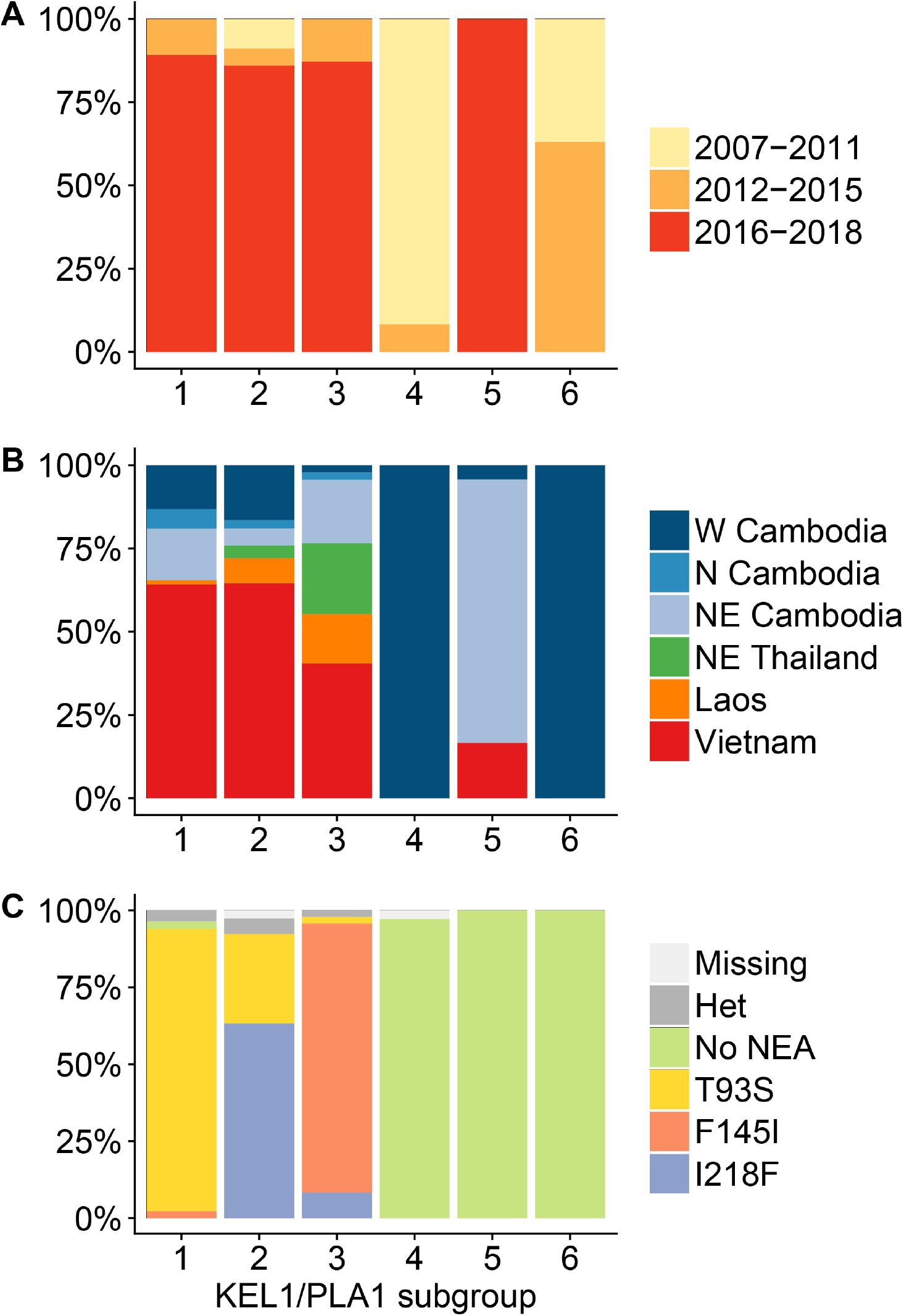
Distinct epideim ological and genetic properties of KEL1/PLA1 subgroups. (A-B) Sample proportions by sampling period (A) and location (B) in the six largest groups of high-similarity KEL1/PLA1 parasites (together comprising >50% of KEL1/PLA1 samples). Subgroups 1-3 emerged recently and are internationally distributed, while subgroups 4 and 6 are older and confined to Western Cambodia. (C) CRT haplotypes in the same groups: newly emerging CRT mutations are highly prevalent in the newer subgroups 1-3, but absent from the older geographically restricted subgroups 4 and 6, and also in subgroup 5, which has recently expanded in northeast Cambodia.

### Emerging CRT mutations associated with KEL1/PLA1 subgroup expansions

The recent, rapid international expansion of some KEL1/PLA1 subgroups after several years of confinement in Cambodia raises the question of whether new genetic changes have produced advantageous phenotypic effects in these subgroups. One candidate set of driver mutations are substitutions in the *crt* gene (PF3D7_0709000) that were recently associated with piperaquine resistance^22^. To investigate this possibility, we calculated the changes in allele frequency in ESEA for all non-synonymous *crt* SNPs from earlier sampling years (2007-2011) to later years (2016-2018) (Supplementary Table 3). Three of these mutations (T93S, F145I and I218F), here referred to as *Newly Emerging Alleles* (NEAs), were rare in 2007-2011 (frequency ≤1%) and rose in frequency by >5% by 2016-2018 (24%, 7% and 13% of all ESEA samples, respectively). These mutations were mutually exclusive: in the entire analysis dataset, we did not identify any sample with multiple NEAs, despite their being present in the same geographic regions in the same time periods.

We found that NEAs occurred on top of a specific constellation of other, more prevalent *crt* mutations (Supplementary Figure 8). First, this background comprised K76T and several other *crt* mutations in common with all chloroquine resistant (CQ-R) parasites^24^. Second, NEAs were only found on the main CQ-R haplotype in ESEA, denoted CVIET (named by *crt* amino acid positions 72-76). Third, they occurred mainly on top of *crt* mutations N326S and I356T, which have previously been associated with artemisinin resistant *kelch13* variants^11^. Finally, NEAs were mainly (though not exclusively) found in KEL1/PLA1 parasites. Four other mutually exclusive *crt* alleles found at lower frequencies than the NEAs (H97Y, H97L, M343I and G353V) were also associated with the same genetic background, with the exception of H97L, which occurred only in non-PLA1 parasites. Consistent with the spread of KEL1/PLA1 and rising frequency of NEAs, SNPs associated with the CVIET haplotype have all increased in frequency over the study period at the expense of those in other *crt* haplotypes (Supplementary Table 3).

NEAs were embedded within long shared haplotypes, with reduced genetic diversity across the whole of chromosome 7 (Supplementary Figures 9-11). This denotes limited breakdown through recombination, typical of a very recent selective sweep. Consistent with their recent origin, NEAs were mainly found in newer KEL1/PLA1 outbreak subgroups: T93S and F145I were almost at fixation in subgroups 1 and 3, respectively, while subgroup 2 contained a mixture of T93S and I218F parasites (Figure 4C, Supplementary Figure 6). In summary, our data suggest that multiple KEL1/PLA1 genetic subgroups were able to spread rapidly across borders in separate transmission waves following the acquisition of one of several mutually exclusive *crt* mutations, which have emerged on a complex genetic background comprising a constellation of other *crt* mutations that have accumulated over decades in ESEA.

Northeast Thailand provides a case study in the genomic epidemiology of these spreading multidrug resistant parasites. In 2011, all sampled parasites from northeast Thailand were *kelch13* R539T mutants, and possessed neither the *plasmepsin*-*II*/*III* amplification nor any NEAs. Although they had delayed parasite clearance times^9,11,16^, they were sensitive to piperaquine, such that DHA-PPQ remained an effective treatment. These parasites had exceptionally low genetic diversity (Supplementary Figure 3), perhaps reflecting population collapse due to effective malaria control efforts. By 2017, however, KEL1/PLA1 had entirely replaced the previous R539T population, with a corresponding rise in DHA-PPQ resistance and treatment failure rates. While the majority of these parasites were from subgroup 3 and possessed the *crt* F145I mutation, we found other NEAs in this area, suggesting that multiple “enhanced” KEL1/PLA1 subgroups, possessing distinct and mutually exclusive *crt* mutations, have independently invaded northeast Thailand, replacing earlier parasite populations.

## Discussion

After a decade of progress^25^, malaria incidence and mortality have been increasing since 2015, prompting the World Health Organisation (WHO) to warn that “without urgent action, we risk going backwards, and missing the global malaria targets for 2020 and beyond”^26^. Major challenges to achieving elimination targets include inadequate funding, parasite drug resistance and insecticide resistance in mosquito vectors^27^. Previous work described a worrying situation unfolding in Southeast Asia over the period 2007-2013, with the emergence of a dominant parasite co-lineage, KEL1/PLA1, that spread across Cambodia causing DHA-PPQ treatment failure^13^. Here, we describe the ongoing evolution and expansion of multidrug resistant *P. falciparum* using whole genomes sampled across ESEA collected up to early 2018.

Our data clearly demonstrate that KEL1/PLA1 has continued its spread out from western Cambodia and is now highly prevalent in multiple regions including northeast Thailand, Vietnam and south Laos, where it has frequently replaced previous indigenous populations. Regardless of the sampling location, KEL1/PLA1 parasites were genetically distinct from non-KEL1/PLA1 parasites, reflecting their recent shared ancestry from western Cambodia. The genomic data show that underlying this spread is not a single lineage but multiple subgroups of KEL1/PLA1 parasites, which have spread widely across ESEA in independent transmission waves. These recently expanding subgroups carry newly emerging alleles (NEAs) in the *crt* gene, which have arisen on a specific constellation of background *crt* mutations and most frequently in KEL1/PLA1 parasites. The rapid rise in frequency of these *crt* alleles suggests that they are markers of an advantageous phenotype. One NEA (F145I) has been shown to reduce piperaquine sensitivity *in vitro*^22^, and a new clinical study (van der Pluijm *et al.* in this issue) demonstrates that NEAs are associated with a higher rate of DHA-PPQ treatment failures. Other *crt* alleles arising on a similar genetic background may also be functionally significant – G353V and H97Y have been associated with reduced piperaquine sensitivity *in vitro*^22^ and H97Y with increased DHA-PPQ treatment failures (van der Pluijm *et al.*). Thus, several novel *crt* variants may be capable of reducing parasite sensitivity to piperaquine, and among these the three NEAs are the alleles whose recent rise in frequency is most conspicuous in our dataset.

KEL1/PLA1 parasites emerged in western Cambodia and expanded within Cambodia during an initial phase starting around 2008. They progressively replaced the local parasite population, such that by 2014 nearly all parasites sampled from western Cambodia were KEL1/PLA1.^13^ This was likely driven by DHA-PPQ resistance, consistent with the association found between *plasmepsin*-*II*/*III* amplification and piperaquine resistance in parasites collected before NEAs were found at significant frequencies^17,18^. After several years of continued exposure to DHA-PPQ, Cambodian parasites acquired further mutations, notably in the *crt* gene, and this may have produced an enhanced phenotype conferring higher-level DHA-PPQ resistance. KEL1/PLA1 subgroups possessing these *crt* mutations were then able to spread rapidly into surrounding regions in the 2015-2018 period.

These findings illustrate an evolutionary process in action. Artemisinin resistance first began as delayed parasite clearance caused by many mutually exclusive *kelch13* mutations, which were generally at low frequency and restricted in geographic range. Over time, a single *kelch13* mutation (C580Y) has become dominant in ESEA, in association with several mutations such as variants in *ferredoxin*, *arps10*, *mdr2* and *crt*^11,12^. Thus, a *soft sweep* (many different, independently emerging advantageous *kelch13* mutations) turned into a *hard sweep* of the *kelch13* C580Y variant as the KEL1/PLA1 co-lineage, resistant to both artemisinin and piperaquine, rose in frequency and swept through Cambodia^13^ and beyond. Following that harder sweep, the parasites have diversified into separate evolutionary branches with emerging new properties. This perhaps reflects a general tendency for diversification after a hard sweep, as the population “explores” a vast evolutionary space and acquires new mutations. The genetic background that has accumulated in ESEA appears to underpin the emergence of *crt* NEAs, as multiple alleles have arisen independently within the last few years that are absent elsewhere. It remains to be seen whether one of these NEAs will becomes dominant and drive a new hard sweep, as *kelch13* C580Y did.

The spread of KEL1/PLA1 up to 2013 described by Amato *et al.*^13^ has worsened significantly. By analogy with cancer biology, KEL1/PLA1 can be viewed as an aggressive cell line that, since 2013, has metastasised, invading new territories and acquiring new genetic properties. This emphasises the importance of surveillance systems in guiding and accelerating malaria elimination. It is possible that prolonged use of DHA-PPQ in Cambodia after resistance to these agents first emerged created the selective pressure for the evolution of enhanced KEL1/PLA1 subgroups. To support national malaria control programmes in their decisions on first-line therapies, particularly given the spread and intensification of resistance, requires genetic surveillance and effective translation of its results. It is with this intent that a large number of the recent samples analysed in this study, particularly from Vietnam, Laos and Cambodia, were contributed by national malaria control programmes collaborating with GenRe-Mekong, a genetic surveillance project operating across the Greater Mekong Subregion. It is hoped that such surveillance tools will support effective elimination of highly drug resistant *P. falciparum* from the Greater Mekong sub-region, and control and elimination efforts elsewhere.

## Materials and Methods

### *P. falciparum* whole genome sequence data

We analysed whole genome sequence data from samples included in the MalariaGEN *P. falciparum* Community Project (https://www.malariagen.net/projects/p-falciparum-community-project) Pf6.2 data release. A large proportion of the samples were collected in clinical observational or drug efficacy studies as detailed in previous publications^1,9,12,13^. Recently collected, previously unpublished samples were contributed by two large-scale multi-site projects: the Tracking Artemisinin Resistance Collaboration II (TRAC2), and the Genetic Reconnaissance in the Greater Mekong Subregion project (GenRe-Mekong, https://www.malariagen.net/projects/spotmalaria). The TRAC2 study (see van de Pluijm *et al.* in this issue) conducted detailed clinical drug efficacy trials at seven sites in the ESEA region, contributing DNA from both dried blood spots (DBS) and leukocyte-depleted venous blood samples. Up to 120 symptomatic patients per site were enrolled, all of whom provided written informed consent under protocols approved by the relevant local ethics authorities and by the Oxford Tropical Research Ethics Committee (OxTREC), Oxford, UK. The GenRe-Mekong project contributed samples from three genetic surveillance projects in partnership with malaria control programmes in Vietnam (IMPE-QN: Institute of Malariology, Parasitology and Entomology Quy Nhon), Lao PDR (CMPE: Centre of Malariology, Parasitology, and Entomology) and Cambodia (Center for Parasitology, Entomology, and Malaria Control). DBS samples were collected from RDT-positive symptomatic patients on admission at provincial hospitals and district health centres in multiple provinces of each country (Supplementary Table 1). All patients provided informed consent (oral in the Lao PDR, written in Vietnam and Cambodia) under protocols approved by the relevant local ethics authorities and OxTREC. These surveillance projects did not involve any clinical component, and no clinical or personal patient data were used in this analysis. The MalariaGEN Analytics website (www.malariagen.net/analytics) provides further details on participating studies and field sites, sample processing and whole genome sequencing approach.

### Whole genome sequencing and dataset selection

DNA from DBS samples underwent selective whole genome amplification^28^ prior to sequencing. Sequence data were generated at the Wellcome Sanger Institute with Illumina short-read technology, and read counts for genotyped variants were called with a standardised analysis pipeline^29^. Samples were genotyped at 1,043,334 biallelic SNPs in the nuclear genome, which were discovered and quality-filtered in the Pf6.0 release (*manuscript under preparation*). Genotypes were called only with a coverage ≥5 reads, and alleles were disregarded when represented by fewer than 2 reads, or 5% of reads when coverage was >50.

To minimize errors and biases, we excluded from analysis all known or suspected duplicate samples, samples from time sequences and from recurrences, samples sequenced with reads <75 nucleotides, and those that had insufficient coverage at >25% of the SNPs. We removed all SNPs that either were invariant or had insufficient coverage in >25% of the remaining samples, leaving 56,026 SNPs to be used in our analysis. We used genotypes at these SNPs to estimate *F*_*WS*_ as previously described ^29^, and removed samples with *F*_*WS*_<0.95, yielding a final set of 1,673 essentially monoclonal samples to be analysed.

### KEL1, PLA1 and *crt* haplotype classification

To identify mutations in the *P falciparum kelch13* gene, we scanned all sequencing reads that align to amino acid positions >350 in this gene, listing any nonsynonymous variants found. Samples where no nonsynonymous mutations were found were listed were labelled as wild-type, unless >25% of position had insufficient coverage for genotyping, in which case the sample was labelled as *undetermined* for *kelch13*. The remaining samples were labelled according to the kelch13 mutation found, or as *heterozygous* if the mutation site was heterozygous, or if more than one nonsynonymous variant was found in the sample. For our analysis of C580Y mutants, we only selected samples labelled for that mutation, disregarding heterozygous samples as these may contain a proportion of non-C580Y parasites.

To assign membership to the KEL1 lineage, we tested the five SNPs characteristic of this haplotype^13^. Moving away from the *kelch13* gene and ignoring missing genotypes, we counted how many of these positions carried a KEL1 characteristic allele, before encountering a position with the other allele. Samples with a total of three or more characteristic allele were labelled as KEL1. PLA1 parasites were identified by scanned all sequencing reads for the characteristic duplication breakpoint^13^. The *crt* haplotypes were determined from the genotypes at the positions that characterize the haplotypes (see Results). A sample was assigned a given haplotype if the corresponding position was mutated while the remaining five had wild type alleles (with the exception of the *crt*218F characteristic position, which has relatively high missingness and was ignored when missing in the presence of another characteristic mutation). The remaining samples were assigned to the “no *crt*” group if all six positions were found to be wild-type, and “missing” if the genotype could not be called at all the mutations.

### Population Structure Analysis

Analyses were performed using a combination of custom software programs written in Java, R and Python using the toolkit *Scikit*-*Allel* (https://scikit-allel.readthedocs.io/en/latest/). To study population structure, we constructed an *NxN* pairwise distance matrix, where *N* is the number of samples, using a previously published procedure ^12^. Analyses of relatedness were conducted using the R language, and specifically the `hclust` hierarchical clustering function and the `cmdscale` implementation of the Classical Multidimensional Scaling method for PCoA.

## Acknowledgements

This study was funded by the Wellcome Trust (098051, 206194, 090770, 204911, 106698/B/14/Z), Bill & Melinda Gates Foundation (OPP1040463, OPP11188166), the Medical Research Council (G0600718), the UK Department for International Development (201900, M006212), and the Intramural Research Program of the National Institute of Allergy and Infectious Diseases. WLH is funded by Cambridge University Hospitals NHS Foundation Trust and National Institute for Health Research. The funders of the study had no role in study design, data collection, data analysis, data interpretation, or writing of the report. This study used data from the MalariaGEN Pf3k Project and Plasmodium falciparum Community Project. Genome sequencing was done by the Wellcome Sanger Institute (WSI), and sample collections were coordinated by the MalariaGEN Resource Centre. We thank the staff of the WSI Sample Logistics, Sequencing, and Informatics facilities for their contribution; Mihir Kekre and Katie Love for their support in the sample processing pipeline; Nick Harding, Alistair Miles, Chris Clarkson and Jacob Almagro-Garcia at WSI and the Oxford Big Data Institute for their helpful suggestions on bioinformatic analyses; and Victoria Cornelius and Christa Henrichs for their support in coordinating and managing the project and the MalariaGEN Plasmodium falciparum Community Project. We thank all patients and collaborators contributing samples to the MalariaGEN Plasmodium falciparum Community Project, in particular investigators who submitted samples used in this study: Chris Plowe, Pascal Ringwald, Elizabeth Ashley, Xinxhuan Su, Maciej Boni. We also extend our thanks to collaborators to the GenRe-Mekong Project: Pannapat Masingboon, Sonexay Phalivon, Nguyen Kim-Tuyen, Thanat Chookajorn, Namfon Kotanan, Phrutsamon Wongnak, Zoë Doran, Salwaluk Panapipat, Ipsita Sinha, Sara Canavati.

## SUPPLEMENTARY MATERIALS

### Supplementary tables

**Supplementary Table 1.** Counts of samples analysed for the different ESEA regions surveyed.

| Region |  |  | Sample<br>Count |
| --- | --- | --- | --- |
| Symbol | Name | Provinces |  |
| WKH | W Cambodia | Pailin, Pursat, Battambang | 518 |
| NKH | N Cambodia | Preah Vihear | 126 |
| NEKH | NE Cambodia | Stung Treng, Ratanakiri | 280 |
| NETH | NE Thailand | Sisaket | 41 |
| LA | Lao PDR | Savannakhet, Salavan, Sekong, Attapeu, Champasak | 250 |
| VN | Vietnam | Binh Phuoc, Dak Lak, Dak Nong, Gia Lai, Quang Tri, Ninh Thuan, Khanh Hoa | 458 |
| Total |  |  | 1673 |

**Supplementary Table 2.** Characteristics of identified KEL1/PLA1 subgroups. For each of the subgroups containing at least 5 samples, we show its distribution by region (left), year of sampling (middle) and *crt* haplotype (right). Parasites classified as “*undetermined*” in the latter section had insufficient coverage of the *crt* gene to determine their haplotype reliably or had ambiguous (het) calls.

| Subgroup | Region |  |  |  |  |  | Collection year |  |  | crt haplotype |  |  |  |  |
| --- | --- | --- | --- | --- | --- | --- | --- | --- | --- | --- | --- | --- | --- | --- |
|  | WKH | NKH | NEKH | NETH | LA | VN | 2007-2011 | 2012-2015 | 2016-2018 | T93S | F145I | I218F | No NEA | Undetermined |
| 1 | 11 | 5 | 13 | 0 | 1 | 54 | 0 | 9 | 75 | 77 | 2 | 0 | 2 | 3 |
| 2 | 13 | 2 | 4 | 3 | 6 | 51 | 7 | 4 | 68 | 23 | 0 | 50 | 0 | 6 |
| 3 | 1 | 1 | 9 | 10 | 7 | 19 | 0 | 6 | 41 | 1 | 41 | 4 | 0 | 1 |
| 4 | 36 | 0 | 0 | 0 | 0 | 0 | 33 | 3 | 0 | 0 | 0 | 0 | 35 | 1 |
| 5 | 1 | 0 | 19 | 0 | 0 | 4 | 0 | 0 | 24 | 0 | 0 | 0 | 24 | 0 |
| 6 | 19 | 0 | 0 | 0 | 0 | 0 | 7 | 12 | 0 | 0 | 0 | 0 | 19 | 0 |
| 7 | 0 | 0 | 9 | 1 | 3 | 4 | 0 | 0 | 17 | 7 | 2 | 0 | 6 | 2 |
| 8 | 0 | 0 | 0 | 0 | 3 | 11 | 0 | 0 | 14 | 13 | 0 | 0 | 0 | 1 |
| 9 | 1 | 0 | 0 | 3 | 8 | 1 | 0 | 2 | 11 | 0 | 0 | 13 | 0 | 0 |
| 10 | 11 | 1 | 0 | 0 | 0 | 0 | 0 | 8 | 4 | 0 | 0 | 0 | 10 | 2 |
| 11 | 9 | 1 | 1 | 1 | 0 | 0 | 1 | 10 | 1 | 0 | 0 | 0 | 12 | 0 |
| 12 | 12 | 0 | 0 | 0 | 0 | 0 | 1 | 6 | 5 | 0 | 0 | 1 | 11 | 0 |
| 13 | 11 | 0 | 0 | 0 | 0 | 0 | 0 | 8 | 3 | 0 | 0 | 0 | 10 | 1 |
| 14 | 7 | 0 | 1 | 2 | 0 | 0 | 0 | 1 | 9 | 0 | 0 | 0 | 10 | 0 |
| 15 | 3 | 5 | 1 | 0 | 0 | 1 | 2 | 4 | 4 | 2 | 0 | 1 | 5 | 2 |
| 16 | 8 | 0 | 0 | 1 | 0 | 0 | 0 | 6 | 3 | 0 | 0 | 0 | 8 | 1 |
| 17 | 1 | 0 | 0 | 0 | 0 | 7 | 0 | 0 | 8 | 2 | 0 | 0 | 6 | 0 |
| 18 | 4 | 3 | 0 | 0 | 0 | 0 | 2 | 5 | 0 | 0 | 0 | 0 | 7 | 0 |
| 19 | 0 | 0 | 5 | 0 | 0 | 0 | 0 | 0 | 5 | 4 | 0 | 0 | 1 | 0 |
| 20 | 5 | 0 | 0 | 0 | 0 | 0 | 4 | 1 | 0 | 0 | 0 | 0 | 5 | 0 |

**Supplementary Table 3.** Variations in frequency *crt* mutations between periods 2007-2013 and 2016-2018. The mutations are grouped by their association to specific genetic backgrounds, i.e. the chloroquine resistant (CQ-R) haplotypes CVIET and CVIDT, and the genetic background on which artemisinin-resistant kelch13 mutations have emerged. The top group (“emerging *crt* mutations”) are those mutations found to increase by at least 1% between the two periods of time in this analysis. Mutations below 1% frequency were disregarded.

| *crt* variant | Frequency in 2007-2011 | Frequency in 2016-2018 | Increase |  |
| --- | --- | --- | --- | --- |
| M74I | 95% | 97% | 1% |  |
| N75E | 77% | 86% | 9% |  |
| K76T | 95% | 97% | 1% |  |
| T93S | 0% | 24% | 24% | NEA |
| H97L | 1% | 6% | 5% |  |
| H97Y | 1% | 2% | 1% |  |
| F145I | 0% | 7% | 7% | NEA |
| I218F | 1% | 14% | 13% | NEA |
| A220S | 95% | 97% | 2% |  |
| Q271E | 95% | 97% | 1% |  |
| N326S | 61% | 76% | 15% |  |
| M343I | 0% | 2% | 2% |  |
| G353V | 1% | 2% | 1% |  |
| I356T | 61% | 76% | 15% |  |
| R371I | 77% | 86% | 9% |  |

### Supplementary figures

**Supplementary Figure 1.**
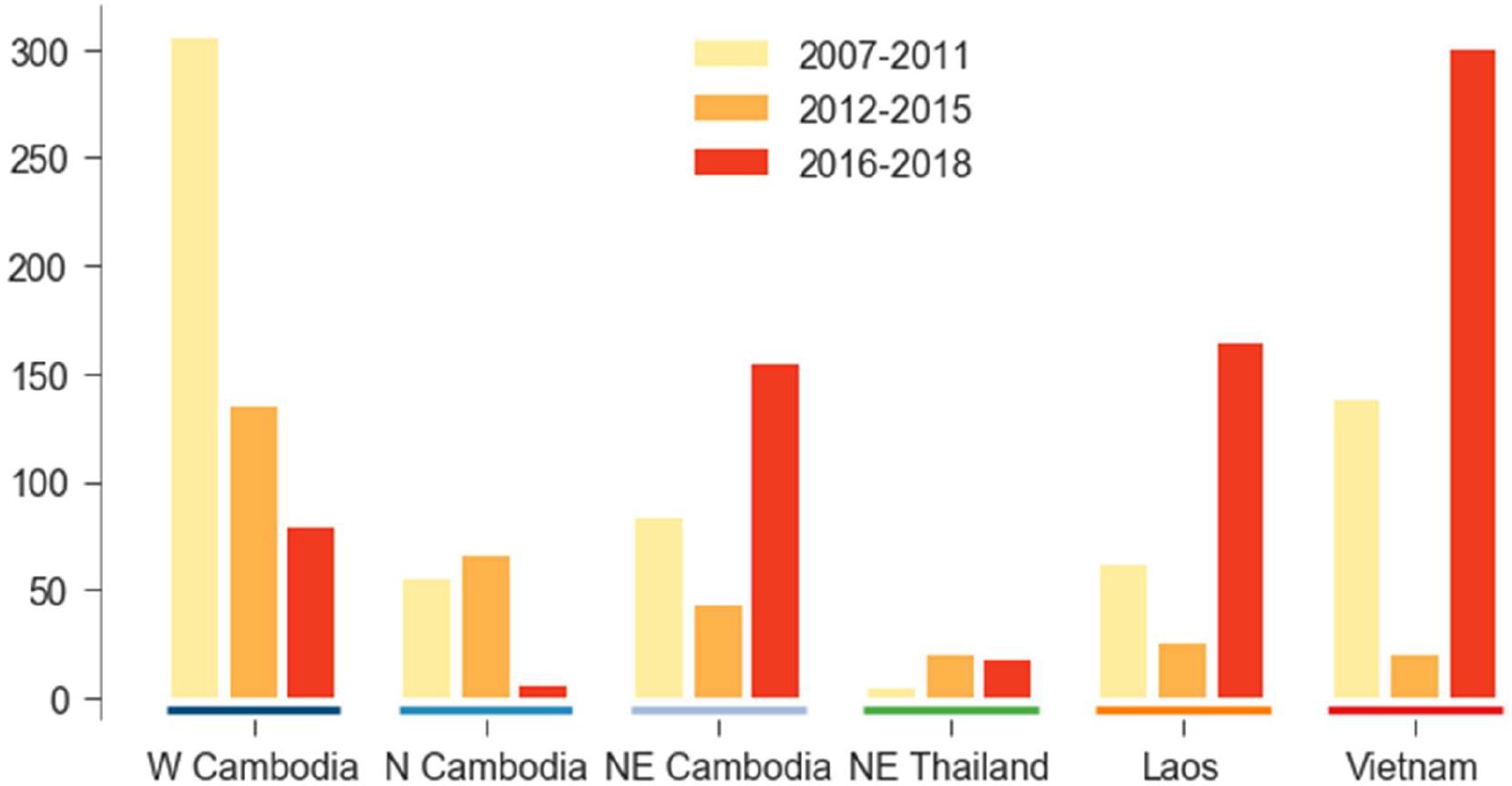
Sample numbers included in main analysis, broken down by country and year. The regions and their symbols are detailed in Supplementary Table 1

**Supplementary Figure 2.**
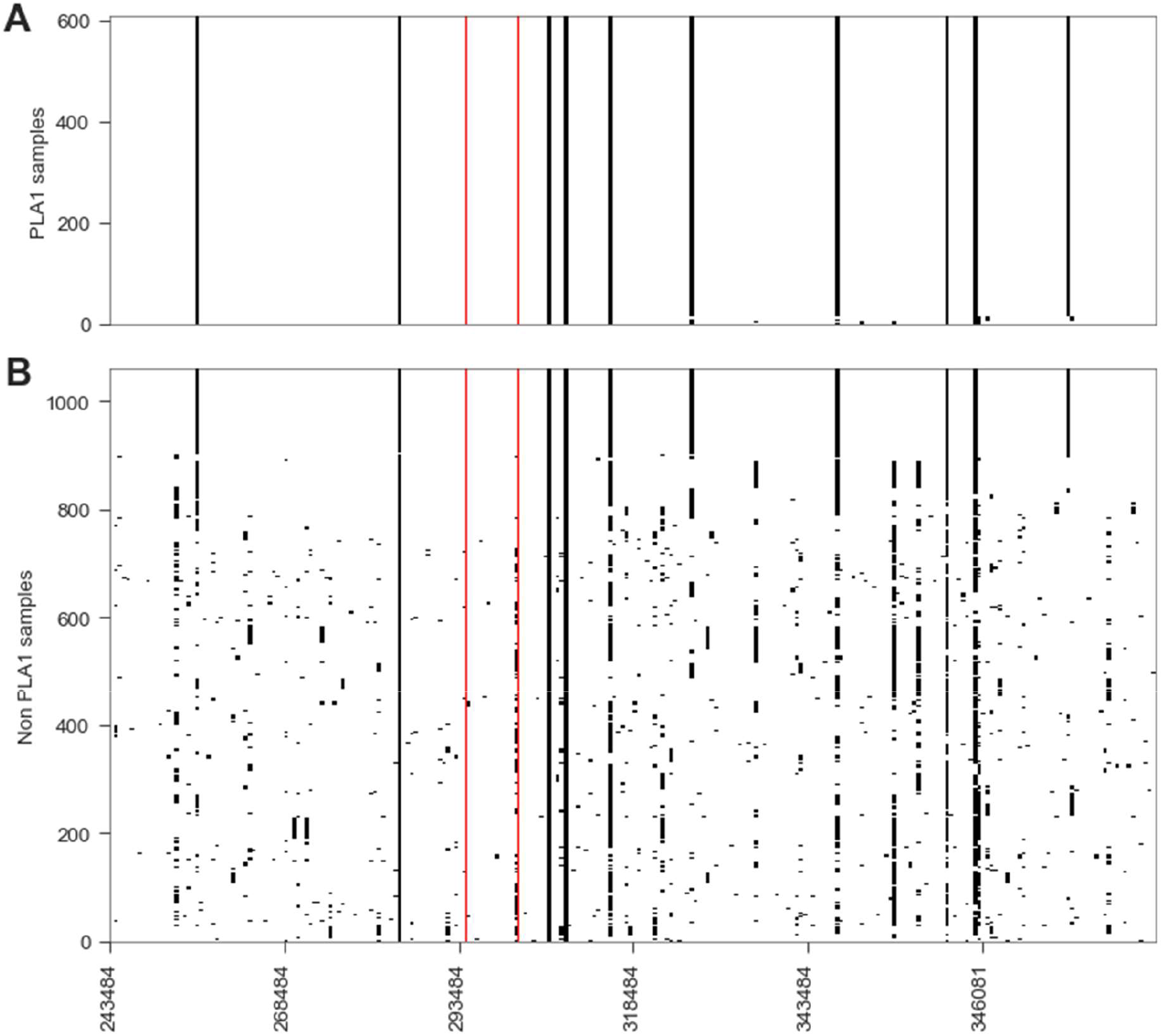
Haplotypes surrounding the *plasmepsin*-*II* and *plasmepsin*-*III* loci for PLA1 samples (A) and non-PLA1 samples (B). Each row represents a sample, and each column represents a single SNP variant. Cells are coloured white for the reference allele (same as 3D7 reference sequence) and black for non-reference allele. The red lines indicate the position of the two *plasmepsin* genes. The presence of a single shared haplotype surrounding these loci is consistent with a single epidemiological origin of the genetic background on which the amplification arose.

**Supplementary Figure 3.**
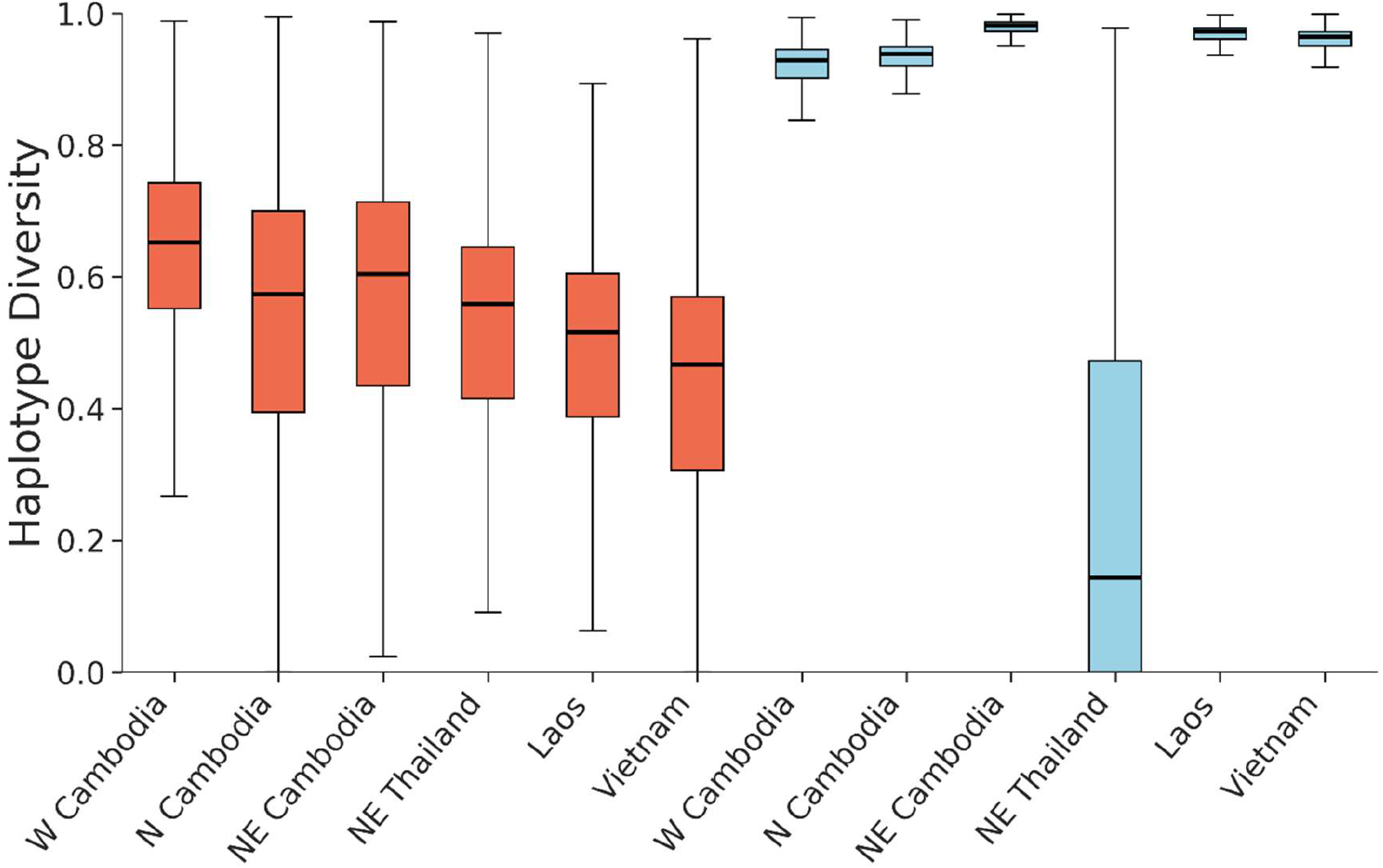
Low haplotype diversity in KEL1/PLA1 parasites. Boxplots show the distribution of haplotype diversity measures in 100-SNP windows along the whole genome for populations of KEL1/PLA1 (red, left) and non-KEL1/PLA1 (blue, right) parasites in the different regions studied.

**Supplementary Figure 4.**
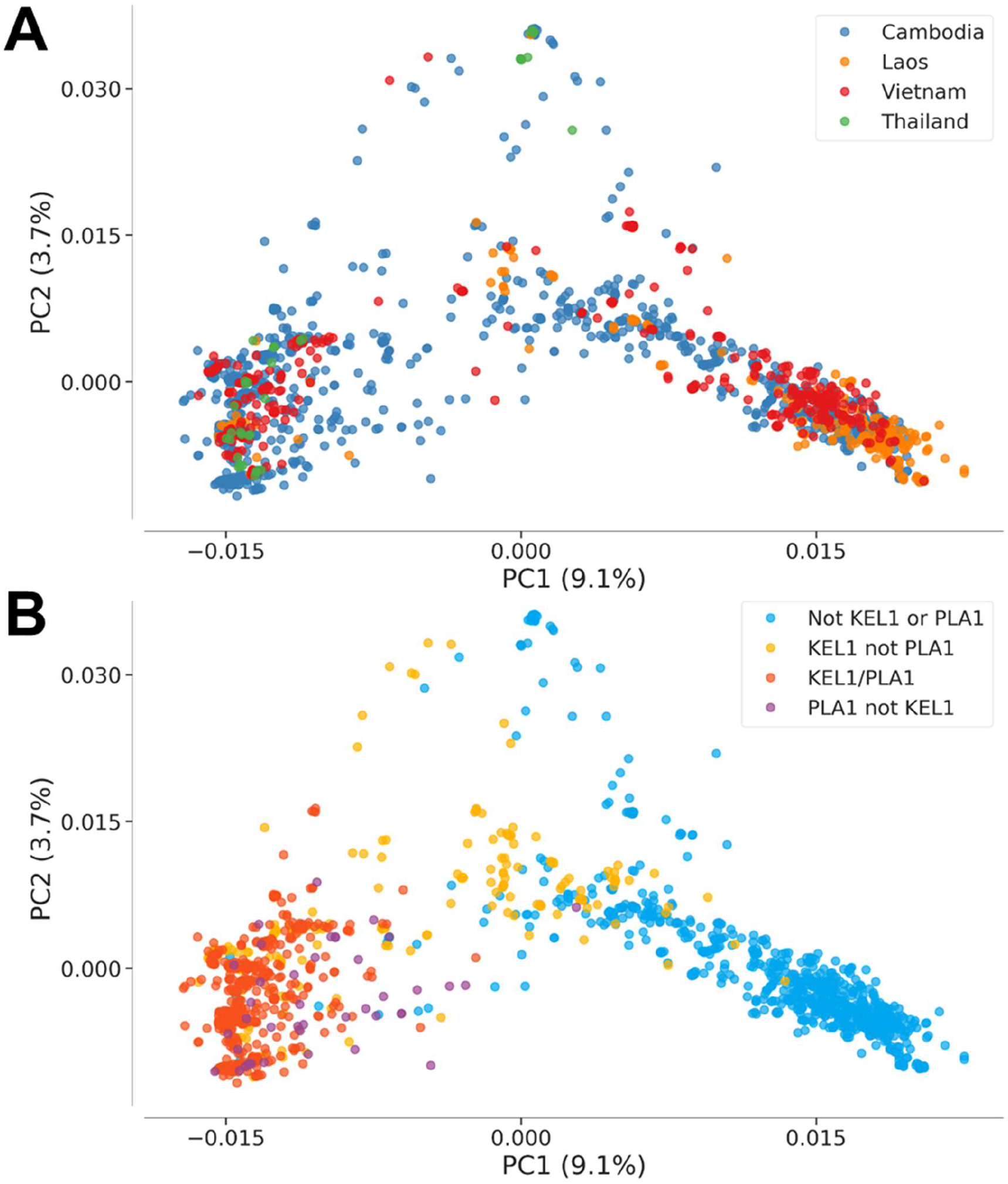
High degree of similarity among KEL1/PLA1 parasites, regardless of geographical origin. Principal Component Analysis (PCoA) based on genetic distance, coloured by country (A) and KEL1/PLA1 status (B). Genetic distances were calculated with correction for linkage disequilibrium and minor allele frequency cutoff of 1%.

**Supplementary Figure 5.**
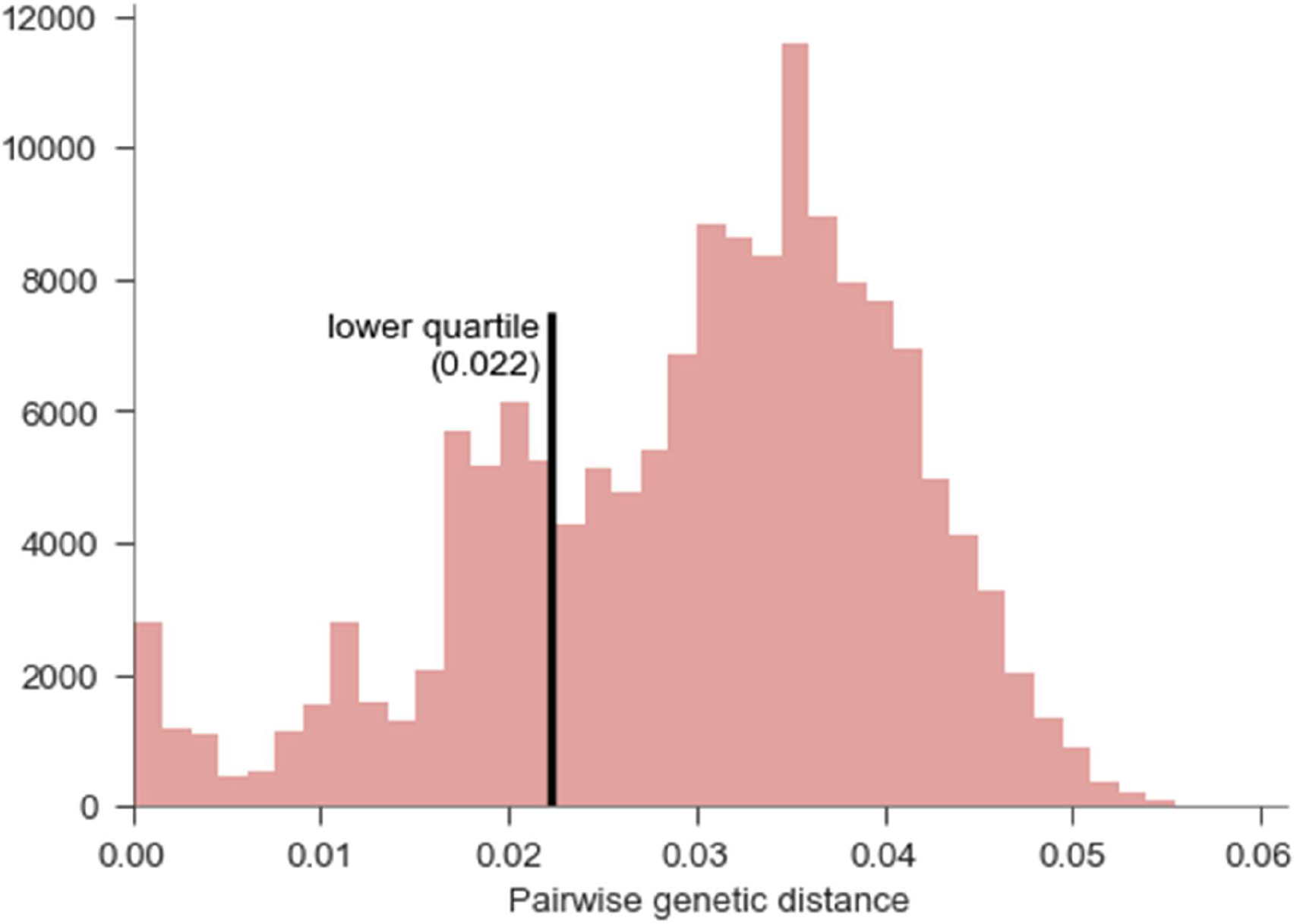
Histogram of pairwise genetic distances for KEL1/PLA1 samples, with lower quartile (used to define related subgroups) marked.

**Supplementary Figure 6.**
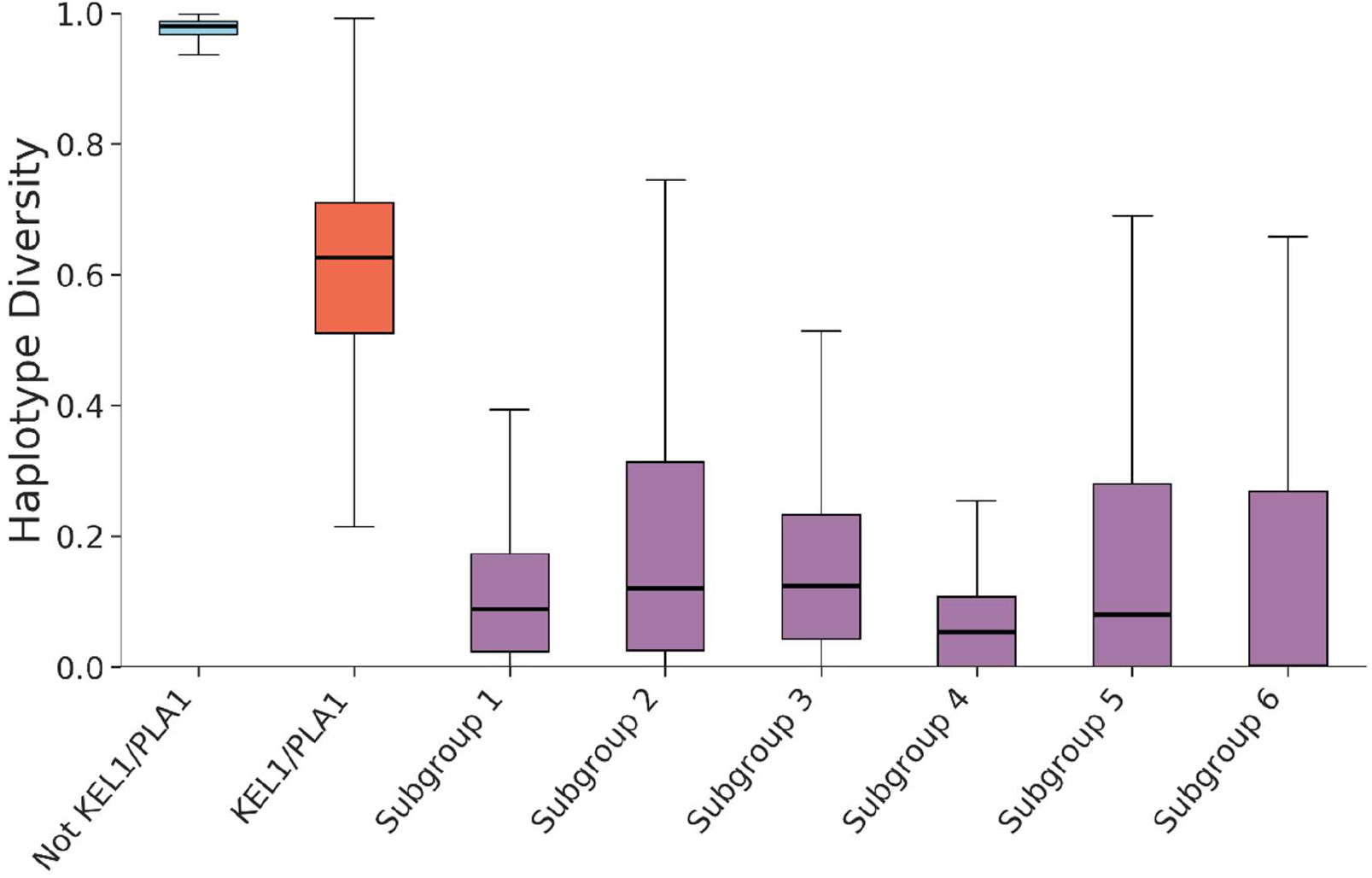
Haplotype diversity in KEL1/PLA1 subgroups. Boxplots show the distribution of aplotype diversity in differnet parasite groups, measured in 100-SNP overlapping windows (50-SNP rolling overlap) across the entire genome.

**Supplementary Figure 7.**
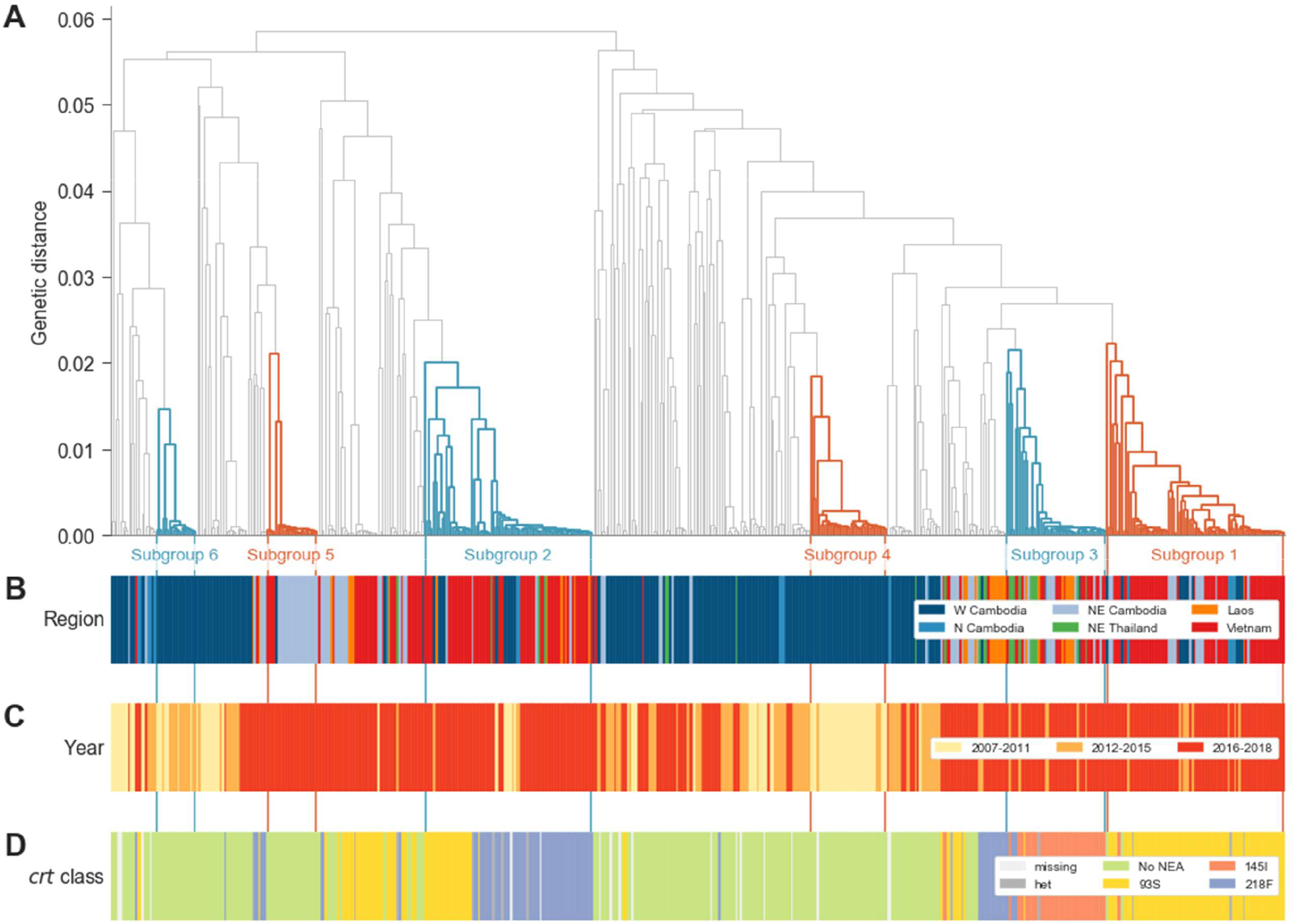
KEL1/PLA1 family tree, geo-temporal distribution and *crt* alleles. (A) Dendrogram of genetic distances within KEL1/PLA1 samples across ESEA, identical to that shown in Figure 3. The six largest subgroups are highlighted and labelled. Colour bars indicate sampling region (B), year (C) and the presence of any Newly Emerging Alleles in *crt*, as defined in main text (D).

**Supplementary Figure 8.**
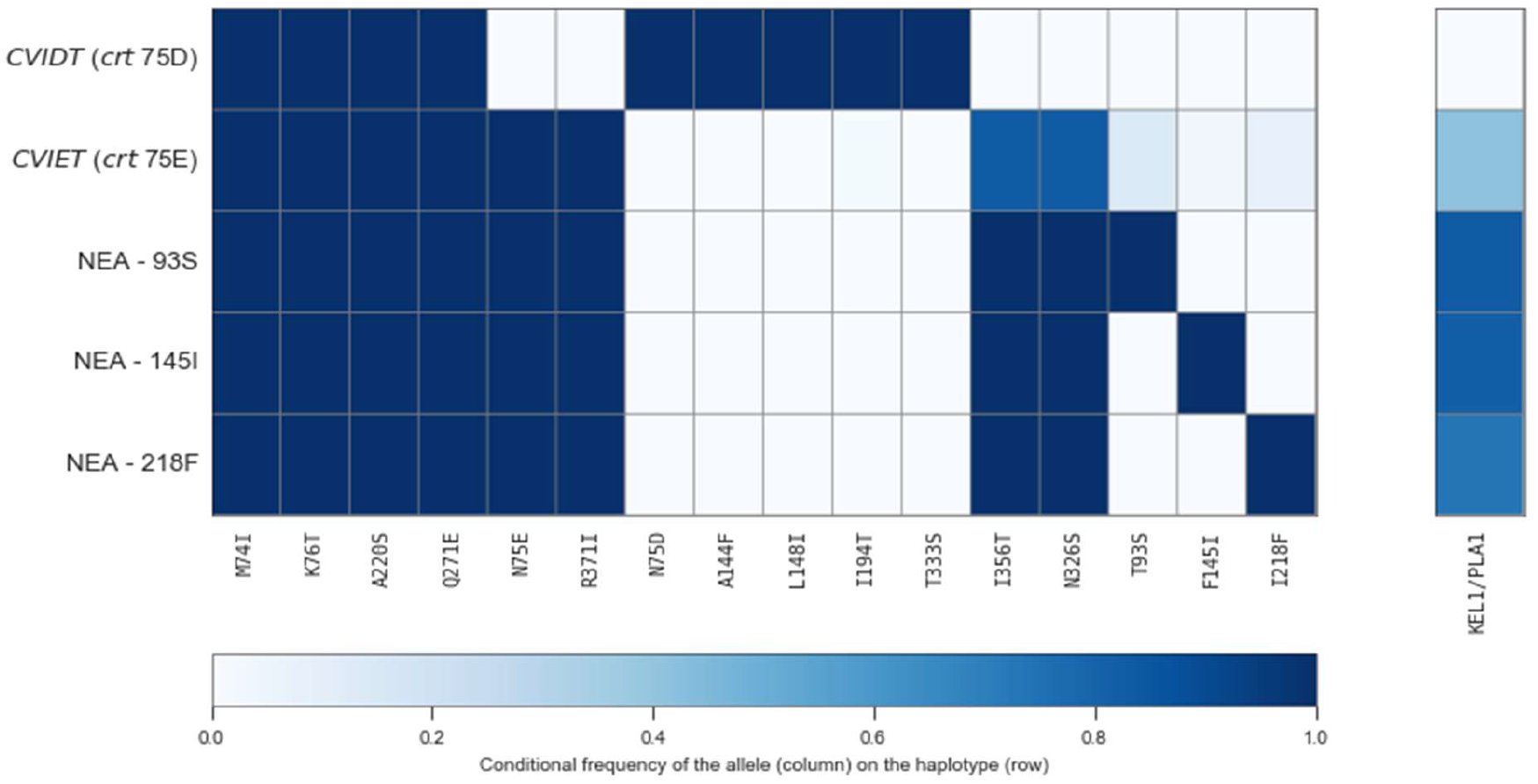
Association of *crt* alleles with specific genetic backgrounds. Each column represents a circulating mutation in the crt gene, and each row represents a genetic background, as mentioned in the main text. Each cell is coloured according to the frequency of the mutation (column) in parasites that carry the specified haplotype (row). NEAs arise on a genetic background comprising the chloroquine resistant CVIET haplotype, the additional mutations at positions 326 and 356, and the KEL1/PLA1 haplotype.

**Supplementary Figure 9.**
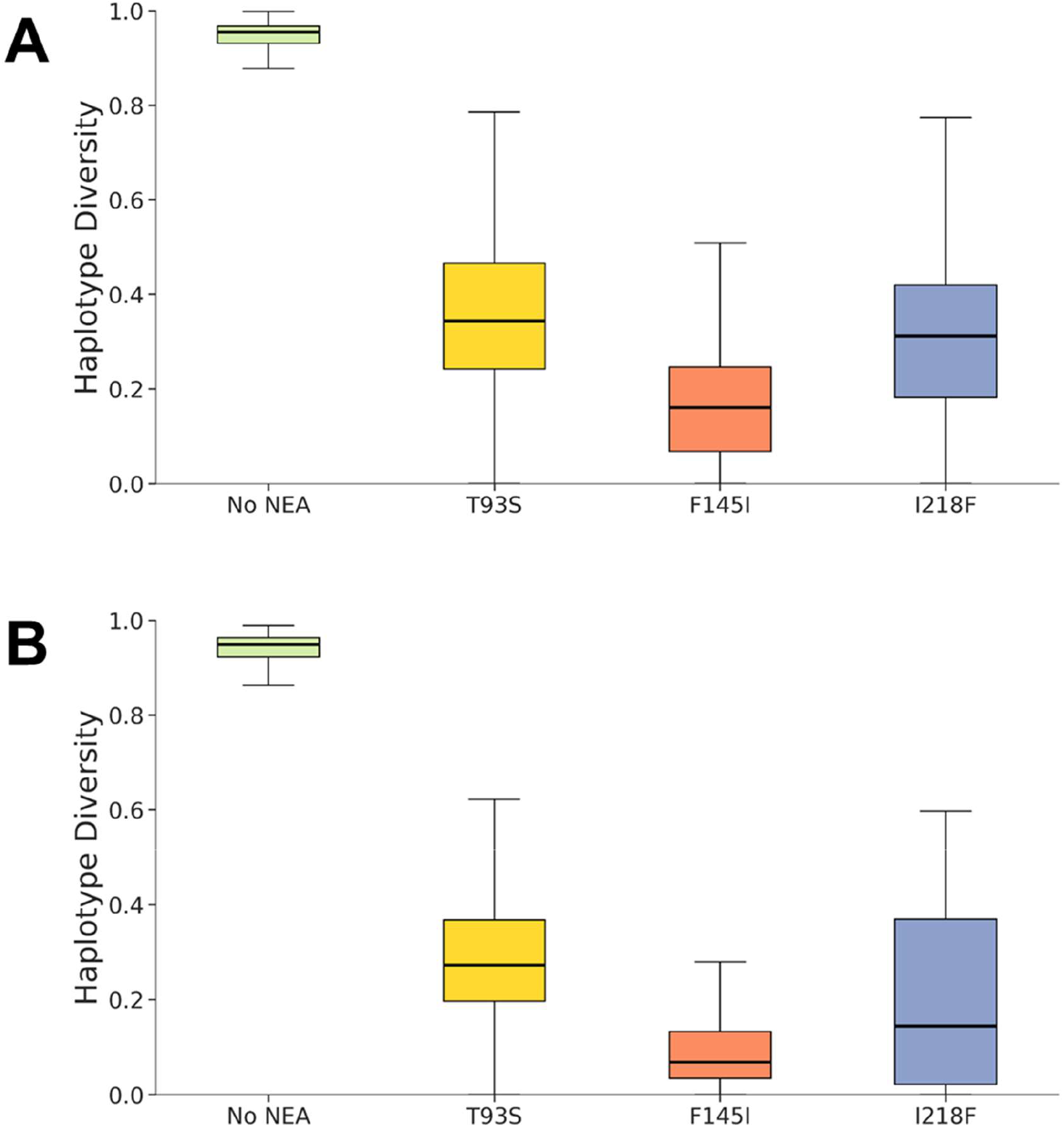
Haplotype diversity across 100-SNP windows in samples possessing Newly Emerging Alleles (NEA) in the crt gene. (A) Distribution of diversity across the whole genome; (B) Distribution of diversity across chromosome 7 only.

**Supplementary Figure 10.**
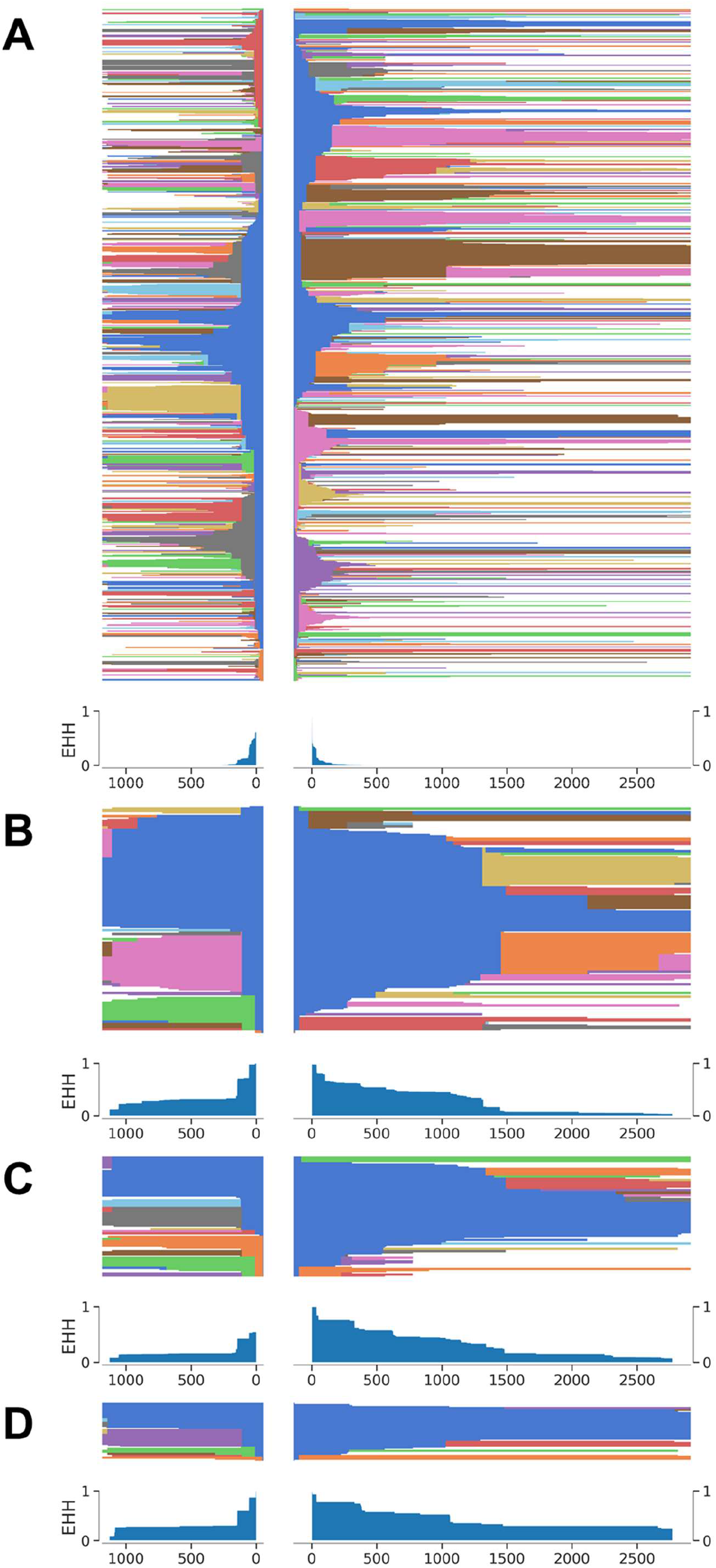
Haplotype frequency plots (above) and extended haplotype homozygosity (EHH, below) along the flanking regions of *crt* on chromosome 7 in samples from each of the following allele groups: no NEA (A), T93S (B), F145I (C) and I218F (D).

**Supplementary Figure 11.**
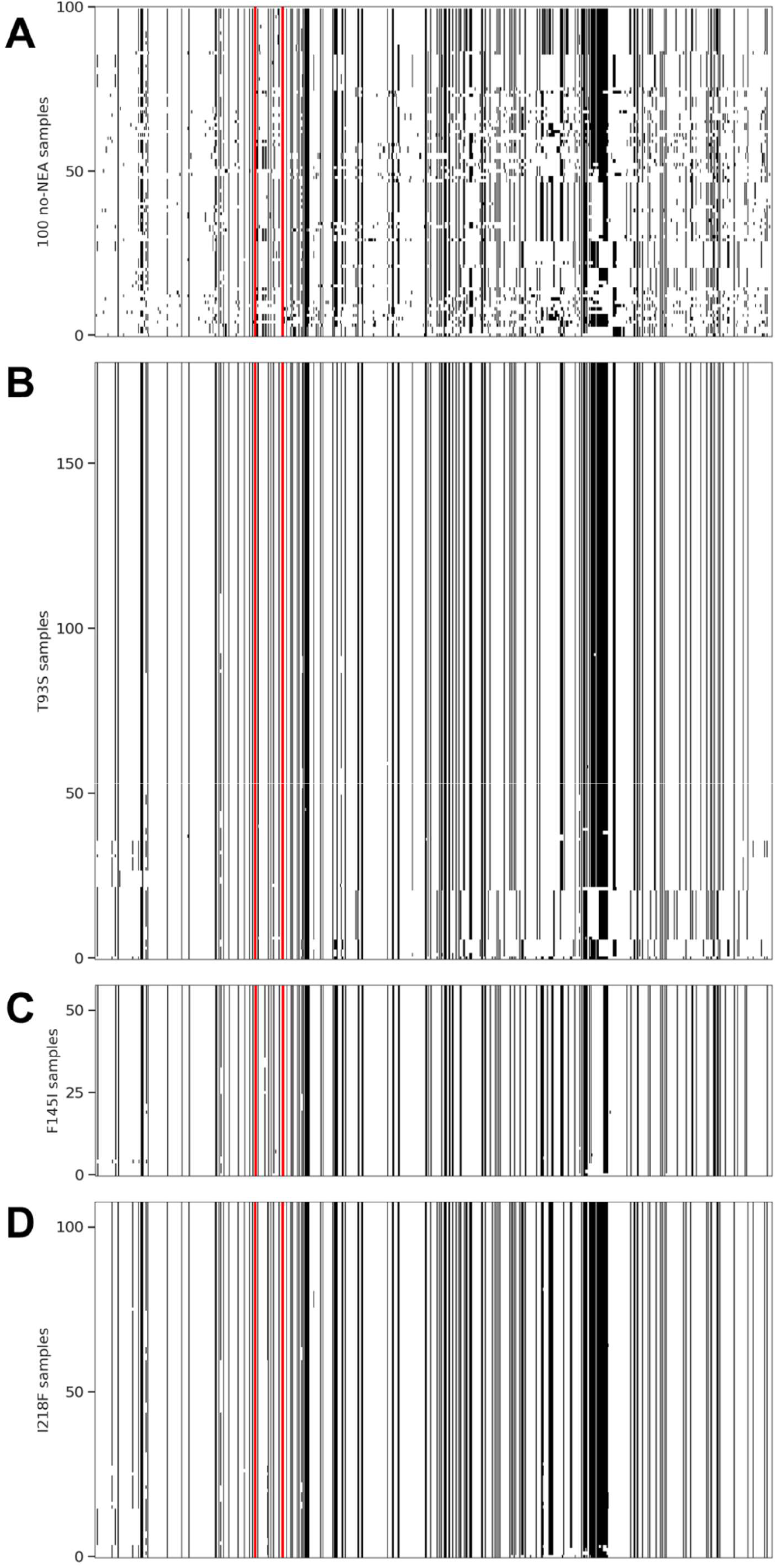
Haplotype across 200KB around *crt* locus on chromosome 7 in 50 randomly selected samples for each of the following allele groups: no NEA (A), T93S (B), F145I (C) and I218F (D).

## References

1. Thanh NV, Thuy-Nhien N, Tuyen NT, et al. Rapid decline in the susceptibility of Plasmodium falciparum to dihydroartemisinin-piperaquine in the south of Vietnam. Malar J 2017; 16(1): 27.

2. Phuc BQ, Rasmussen C, Duong TT, et al. Treatment Failure of Dihydroartemisinin/Piperaquine for Plasmodium falciparum Malaria, Vietnam. Emerg Infect Dis 2017; 23(4): 715–7.

4. Amaratunga C, Lim P, Suon S, et al. Dihydroartemisinin–piperaquine resistance in Plasmodium falciparum malaria in Cambodia: a multisite prospective cohort study. The Lancet Infectious Diseases 2016; 16(3): 357–65.

4. Mita T, Tanabe K, Kita K. Spread and evolution of Plasmodium falciparum drug resistance. Parasitol Int 2009; 58(3): 201–9.

5. Dondorp AM, Nosten F, Yi P, et al. Artemisinin Resistance in Plasmodium falciparum Malaria. N Engl J Med 2009; 361: 455–67.

6. Amaratunga C, Sreng S, Suon S, et al. Artemisinin-resistant Plasmodium falciparum in Pursat province, western Cambodia: a parasite clearance rate study. The Lancet Infectious Diseases 2012; 12(11): 851–8.

7. Phyo AP, Nkhoma S, Stepniewska K, et al. Emergence of artemisinin-resistant malaria on the western border of Thailand: a longitudinal study. The Lancet 2012; 379(9830): 1960–6.

8. Leang R, Barrette A, Bouth DM, et al. Efficacy of dihydroartemisinin-piperaquine for treatment of uncomplicated Plasmodium falciparum and Plasmodium vivax in Cambodia, 2008 to 2010. Antimicrob Agents Chemother 2013; 57(2): 818–26.

9. Ashley EA, Dhorda M, Fairhurst RM, et al. Spread of artemisinin resistance in Plasmodium falciparum malaria. The New England Journal of Medicine 2014; 371(5): 411–23.

10. Miotto O, Almagro-Garcia J, Manske M, et al. Multiple populations of artemisinin-resistant Plasmodium falciparum in Cambodia. Nature genetics 2013; 45(6): 648–55.

11. Miotto O, Amato R, Ashley EA, et al. Genetic architecture of artemisinin-resistant Plasmodium falciparum. Nature genetics 2015; 47(3): 226–34.

12. MalariaGEN Plasmodium falciparum Community Project. Genomic epidemiology of artemisinin resistant malaria. Elife 2016; 5.

13. Amato R, Pearson RD, Almagro-Garcia J, et al. Origins of the current outbreak of multidrug-resistant malaria in southeast Asia: a retrospective genetic study. The Lancet Infectious Diseases 2018; 18(3): 337–45.

14. Imwong M, Hien TT, Thuy-Nhien NT, Dondorp AM, White NJ. Spread of a single multidrug resistant malaria parasite lineage (PfPailin) to Vietnam. The Lancet Infectious Diseases 2017; 17(10): 1022–3.

15. Imwong M, Suwannasin K, Kunasol C, et al. The spread of artemisinin-resistant Plasmodium falciparum in the Greater Mekong subregion: a molecular epidemiology observational study. The Lancet Infectious Diseases 2017; 17(5): 491–7.

16. Ariey F, Witkowski B, Amaratunga C, et al. A molecular marker of artemisinin-resistant Plasmodium falciparum malaria. Nature 2014; 505(7481): 50–5.

17. Amato R, Lim P, Miotto O, et al. Genetic markers associated with dihydroartemisinin–piperaquine failure in Plasmodium falciparum malaria in Cambodia: a genotype–phenotype association study. The Lancet Infectious Diseases 2017; 17(2): 164–73.

18. Witkowski B, Duru V, Khim N, et al. A surrogate marker of piperaquine-resistant Plasmodium falciparum malaria: a phenotype–genotype association study. The Lancet Infectious Diseases 2017; 17(2): 174–83.

19. Bopp S, Magistrado P, Wong W, et al. Plasmepsin II-III copy number accounts for bimodal piperaquine resistance among Cambodian Plasmodium falciparum. Nat Commun 2018; 9(1): 1769.

20. Mukherjee A, Gagnon D, Wirth DF, Richard D. Inactivation of Plasmepsins 2 and 3 Sensitizes Plasmodium falciparum to the Antimalarial Drug Piperaquine. Antimicrob Agents Chemother 2018; 62(4).

21. Agrawal S, Moser KA, Morton L, et al. Association of a Novel Mutation in the Plasmodium falciparum Chloroquine Resistance Transporter With Decreased Piperaquine Sensitivity. J Infect Dis 2017; 216(4): 468–76.

22. Ross LS, Dhingra SK, Mok S, et al. Emerging Southeast Asian PfCRT mutations confer Plasmodium falciparum resistance to the first-line antimalarial piperaquine. Nat Commun 2018; 9(1): 3314.

23. Dhingra SK, Redhi D, Combrinck JM, et al. A Variant PfCRT Isoform Can Contribute to Plasmodium falciparum Resistance to the First-Line Partner Drug Piperaquine. MBio 2017; 8(3).

24. Summers RL, Dave A, Dolstra TJ, et al. Diverse mutational pathways converge on saturable chloroquine transport via the malaria parasite’s chloroquine resistance transporter. Proc Natl Acad Sci U S A 2014; 111(17): E1759–67.

25. Bhatt S, Weiss DJ, Cameron E, et al. The effect of malaria control on Plasmodium falciparum in Africa between 2000 and 2015. Nature 2015; 526(7572): 207–11.

26. WHO. Global response to malaria at crossroads. 2017. http://www.who.int/en/news-room/detail/29-11-2017-global-response-to-malaria-at-crossroads (accessed 01/10/2018 2018).

27. WHO. Key points: World malaria report 2017. 2017. http://www.who.int/malaria/media/world-malaria-report-2017/en/ (accessed 01/10/2018 2018).

28. Oyola SO, Ariani CV, Hamilton WL, et al. Whole genome sequencing of Plasmodium falciparum from dried blood spots using selective whole genome amplification. Malar J 2016; 15(1): 597.

29. Manske M, Miotto O, Campino S, et al. Analysis of Plasmodium falciparum diversity in natural infections by deep sequencing. Nature 2012; 487(7407): 375–9.

